# Improving the phylogenetic resolution of Malaysian and Javan mahseer (Cyprinidae), *Tor tambroides* and *Tor tambra*: Whole mitogenomes sequencing, phylogeny and potential mitogenome markers

**DOI:** 10.1101/2021.04.19.440536

**Authors:** Leonard Whye Kit Lim, Hung Hui Chung, Melinda Mei Lin Lau, Fazimah Aziz, Han Ming Gan

## Abstract

The true mahseer (*Tor* spp.) is one of the highest valued fish in the world due to its high nutritional value and great unique taste. Nevertheless, its morphological characterization and single mitochondrial gene phylogeny in the past had yet to resolve the ambiguity in its taxonomical classification. In this study, we sequenced and assembled 11 complete mahseer mitogenomes collected from Java of Indonesia, Pahang and Terengganu of Peninsular Malaysia as well as Sarawak of East Malaysia. The mitogenome evolutionary relationships among closely related *Tor* spp. samples were investigated based on maximum likelihood phylogenetic tree construction. Compared to the commonly used COX1 gene fragment, the complete COX1, Cytb, ND2, ND4 and ND5 genes appear to be better phylogenetic markers for genetic differentiation at the population level. In addition, a total of six population-specific mitolineage haplotypes were identified among the mahseer samples analyzed, which this offers hints towards its taxonomical landscape.

## Introduction

The *Tor* spp. (true mahseer) is one of the most exorbitant freshwater fish grouped under the Cyprinidae family, valued by their high nutritional value and great food security potential (Day, 1876; Thomas, 1873). These fishes inhabit rapid-flowing waters with rocky bottoms (Shreshtha, 1997). To date, there are a total of 16 *Tor* species identified worldwide and 18.8% (three) of them are found within the freshwaters of Indonesia and Malaysia, namely *Tor tambra, Tor tambroides* and *Tor douronensis* (Ng, 2004).

The environmental degradation within the *Tor* habitat is pacing in an unprecedented elevation trend during the recent years with the increasing human activities like dam construction (Ingram et al., 2005). The IUCN (International Union for Conservation of Nature) Red List assessed three Near Threatened, one Vulnerable and one Critically Endangered *Tor* spp. to date while the others remained Data Deficient (Pinder et al., 2019). The three aforementioned Indonesian-Malaysian *Tor* fish species are among the 11 Data Deficient *Tor* spp. that lack detailed characterization and conservation evaluation. Some studies have also shown that the different colours (silver-bronze and reddish) exhibited in *T. tambroides* may be associated with environmental influences (Esa et al., 2006; Siraj et al., 2007; Vrijenhoek, 1998). Unfortunately, the ambiguous original descriptions of these three *Tor* species had led to misidentification and confusion among the scientific communities in the past (Walton et al, 2017).

A mitochondrial DNA diversity investigation by Walton et al. (2017) and Esa et al. (2008) using the standard cytochrome oxidase I (COX1) gene fragment, had revealed substantial genetic diversity between the *T. tambra* and *T. tambroides* populations sampled from East Malaysia region of the Borneo Island (Batang Ai, Sarawak, East Malaysia), West Java of Indonesia and Peninsular Malaysia (Pahang, Perak, Negeri Sembilan and Kelantan). The utilization of a single gene fragment for analysis is deemed insufficient in terms of data estimates precision and phylogenetic resolution (Duchêne et al., 2011; Walton et al., 2017).

The advent of next-generation sequencing and bioinformatic pipeline enabling the rapid sequencing and assembly of complete mitogenomes from various fish species. Although recent whole mitogenome sequencing of these three *Tor* spp. has shed some lights onto their taxonomical and phylogenetical statuses conservation efforts, the number of publicly available complete mitogenome for *Tor* spp. remains relatively low which precludes detailed mitogenome analyses (Norfatimah et al., 2014; Mohamed Yunus et al., 2018).

In this study, we report a mitogenome-based phylogeny of *Tor tambroides* and *Tor tambra* by sequencing the complete mitogenomes of 11 *Tor* spp. isolated from selected geographical regions across Java and Malaysia. Furthermore, we also identified the SNP markers in all candidate mitochondrial genes that are unique to the identified mitolineages from current taxon sampling effort enabling a more rapid and cost-effective mitochondrial lineage assessment of this fish species.

## Materials and Methods

### Sampling and genomic DNA extraction

Five farmed Sarawak *T. tambroides* fishes were sampled from LTT Aquaculture Farm (1.5482 N 110.5490 E) and Sampadi Fishery Farm (1.5093 N 110.2937 E) whereas the other six fish DNA samples in this study were obtained from Walton et al. (2017) (Table 1 & Supplementary Figure 1). The five farmed fishes were deposited as voucher specimens in the fish museum located at Faculty of Resource Science and Technology, Universiti Malaysia Sarawak. Muscle tissues were collected from these fishes after euthanization in compliance to the guidelines and permit issued by the Animal Ethics Committee of Universiti Malaysia Sarawak (UNIMAS/ TNC(PI)-04.01/06-09 (17). Whole genomic DNA was extracted via the CTAB method as previously described by Chung et al., (2018).

**Table 1.**
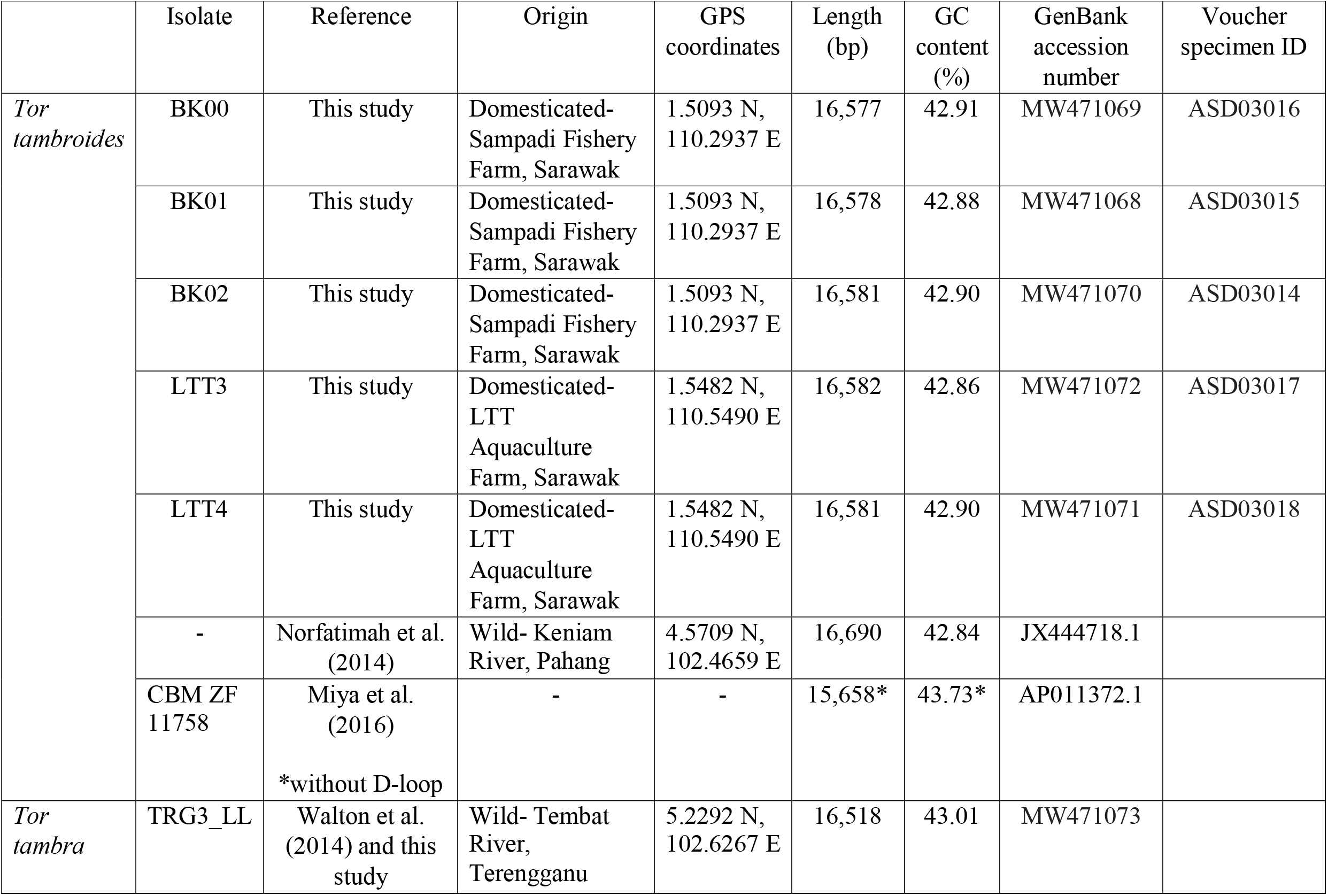
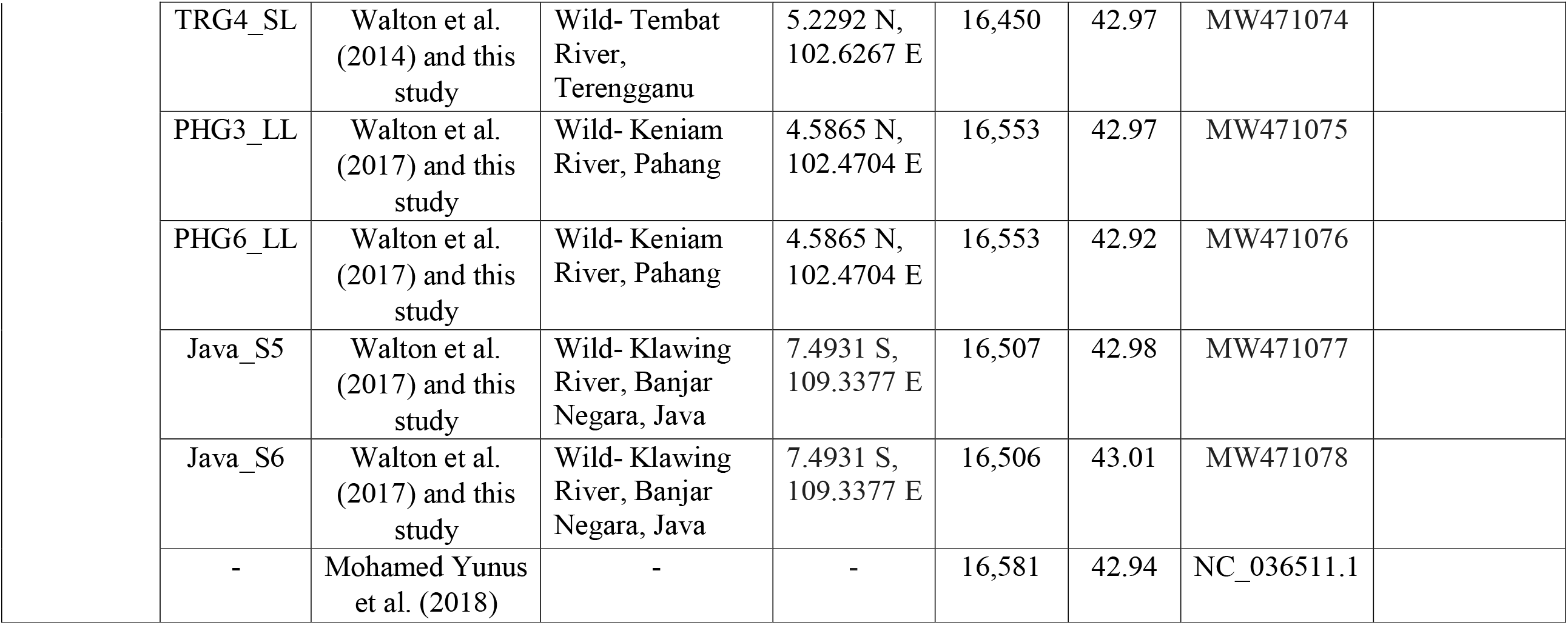
The profiles of all *Tor* spp. fish samples with complete mitogenomes used in this study.

|  | Isolate | Reference | Origin | GPS coordinates | Length (bp) | GC content (%) | GenBank accession number | Voucher specimen ID |
| --- | --- | --- | --- | --- | --- | --- | --- | --- |
| *Tor tambroides* | BK00 | This study | Domesticated-Sampadi Fishery Farm, Sarawak | 1.5093 N, 110.2937 E | 16,577 | 42.91 | MW471069 | ASD03016 |
|  | BK01 | This study | Domesticated-Sampadi Fishery Farm, Sarawak | 1.5093 N, 110.2937 E | 16,578 | 42.88 | MW471068 | ASD03015 |
|  | BK02 | This study | Domesticated-Sampadi Fishery Farm, Sarawak | 1.5093 N, 110.2937 E | 16,581 | 42.90 | MW471070 | ASD03014 |
|  | LTT3 | This study | Domesticated-LTT Aquaculture Farm, Sarawak | 1.5482 N, 110.5490 E | 16,582 | 42.86 | MW471072 | ASD03017 |
|  | LTT4 | This study | Domesticated-LTT Aquaculture Farm, Sarawak | 1.5482 N, 110.5490 E | 16,581 | 42.90 | MW471071 | ASD03018 |
|  | - | Norfatimah et al. (2014) | Wild- Keniam River, Pahang | 4.5709 N, 102.4659 E | 16,690 | 42.84 | JX444718.1 |  |
|  | CBM ZF 11758 | Miya et al. (2016)<br>*without D-loop | - | - | 15,658* | 43.73* | AP011372.1 |  |
| *Tor tambra* | TRG3_LL | Walton et al. (2014) and this study | Wild- Tembat River, Terengganu | 5.2292 N, 102.6267 E | 16,518 | 43.01 | MW471073 |  |

|  |  |  |  |  |  |  |
| --- | --- | --- | --- | --- | --- | --- |
| TRG4_SL | Walton et al.<br>(2014) and this<br>study | Wild- Tembat<br>River,<br>Terengganu | 5.2292 N,<br>102.6267 E | 16,450 | 42.97 | MW471074 |
| PHG3_LL | Walton et al.<br>(2017) and this<br>study | Wild- Keniam<br>River, Pahang | 4.5865 N,<br>102.4704 E | 16,553 | 42.97 | MW471075 |
| PHG6_LL | Walton et al.<br>(2017) and this<br>study | Wild- Keniam<br>River, Pahang | 4.5865 N,<br>102.4704 E | 16,553 | 42.92 | MW471076 |
| Java_S5 | Walton et al.<br>(2017) and this<br>study | Wild- Klawing<br>River, Banjar<br>Negara, Java | 7.4931 S,<br>109.3377 E | 16,507 | 42.98 | MW471077 |
| Java_S6 | Walton et al.<br>(2017) and this<br>study | Wild- Klawing<br>River, Banjar<br>Negara, Java | 7.4931 S,<br>109.3377 E | 16,506 | 43.01 | MW471078 |
| - | Mohamed Yunus<br>et al. (2018) | - | - | 16,581 | 42.94 | NC_036511.1 |

### Mitogenome sequencing and assembly

Approximately 100-500 ng of genomic DNA as quantified by Denovix High Sensitivity Kit was fragmented to 350 bp using a Covaris Ultrasonicator. The fragmented DNA was subsequently prepared using NEB Ultra Illumina Library Preparation kit according to the manufacturer’s instructions. Low coverage sequencing (genome skimming) of the Sarawak mahseers were performed on a NovaSEQ6000 (2 x 150 run configuration). The remaining samples were sequenced on a MiSeq (1 x 150, 2 x 150 or 2 x 250 bp run configuration). The mitogenomes were then assembled using NOVOPlasty version 4.2.1 (Dierckxsens et al., 2017) followed by annotation using MitoAnnotator (Iwasaki et al., 2013). Annotated mitogenome sequences were submitted to GenBank database.

### Phylogenetic analysis

A total of 98 standard 650-bp COX1 gene fragments obtained from the GenBank public database were aligned using MUSCLE version 3.8.31 (Edgar, 2004) and subsequently used to construct maximum likelihood tree with 1000 bootstrap replications via IQ-TREE2 (Nguyen et al., 2015). For mitogenome-based phylogeny, 13 mitochondrial protein coding genes and 2 ribosomal RNA genes from 16 *Tor* fishes (including the 11 *Tor* fishes from this study, two outgroups *Tor tor* and *Tor baraka*, as well as one *T. tambra* and two *T. tambroides* mitogenome sequences available from GenBank) were extracted from the whole mitogenomes of *Tor* spp. and aligned individually using MUSCLE version 3.8.31 (Edgar, 2004). Next, these alignments were concatenated and used for maximum likelihood tree construction as described above.

### Haplotype identification

Haplotypes were detected using DnaSP version 6.12.03 (Rozas et al., 2017) across all the 15 mitochondrial genes (13 protein-coding genes and two ribosomal RNA genes) of 13 *Tor* fishes. The variable nucleotides and positions in each gene were tabulated for each mitolineage-specific haplotype.

## Results and discussions

### Complete *Tor* spp. mitogenome recovery from Illumina-based genome skimming approach

A total of five *T. tambroides* (Sarawak, Malaysia) and six *T. tambra* (Peninsular Malaysia and Java Indonesia) mitogenomes were sequenced in this study (Table 1). The *T. tambroides* fishes originated from Sarawak were farmed fish whereas the *T. tambra* fishes were collected from their natural habitat. The mitogenome sizes of these fishes showed high degree of conservation, especially those yielded from Java and Pahang state (PHG3_LL and PHG6_LL) with subtle length variation in the putative non-coding control region. Among the 11 *Tor* spp. mitogenomes sequenced in this study, the longest mitogenome (16,582 bp) was from the Sarawak state, originating from LTT Aquaculture Farm Sdn Bhd in Sarawak, Malaysia. The other four *T. tambroides* mitogenomes from Sarawak state were longer than that of the *T. tambra* from other localities in this study. The *T. tambra* collected from Terengganu (TRG4_SL) recorded the shortest mitogenome sequences (16,450 bp).

All *Tor* spp. mitogenomes obtained from this study consist of the typical 22 transfer RNA genes, 13 protein coding genes, two ribosomal RNA genes and a non-coding D-loop control region (Figure 1), which echoed the results from the first *T. tambroides* mitogenome sequenced by Norfatimah et al. (2014). The GC contents of *Tor* mitogenomes were also greatly conserved where all of them depicted an AT bias. Interestingly, the *T. tambroides* mitogenomes all have GC contents maintained below 42.92% whereas all the *T. tambra* mitogenomes have GC composition of at least 42.92% in this study. The highest GC contents (43.01%) were discovered in both Java (Java_S6) and Terengganu state (TRG3_LL) *T. tambra* fishes. The *T. tambroides* fish yielded from Sarawak state (LTT3) displayed the lowest GC composition of all (42.86%) in this study.

**Figure 1.**
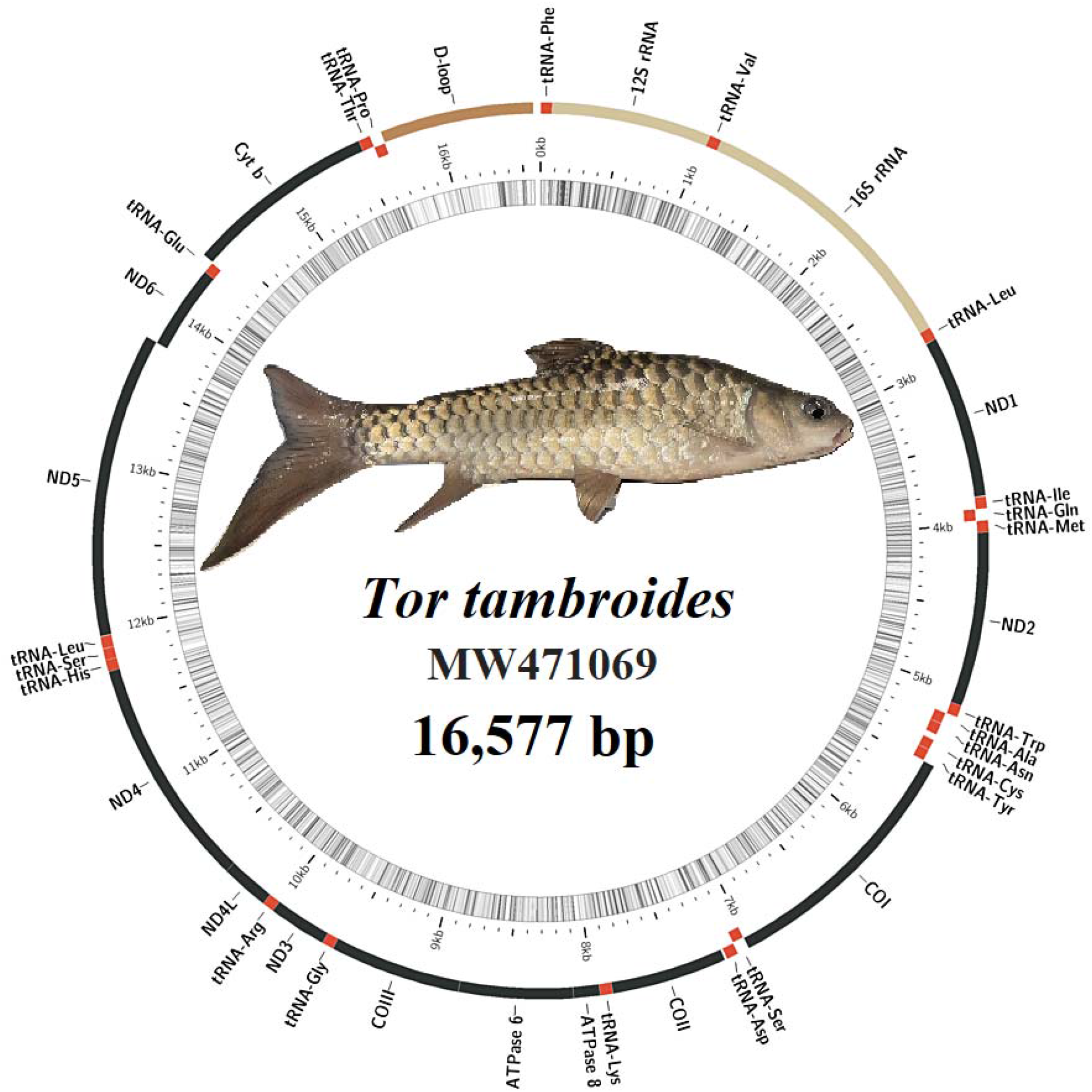
The mitogenome map structure of *Tor tambroides* BK00 (GenBank accession number: MW471069) collected from Sampadi Fishery Farm, Sarawak.

### Standard COX1 gene fragment failed to delineate *Tor* spp. by population

Maximum likelihood phylogenetic tree (Figure 2 & Supplementary Figure 2) constructed based on the alignment of standard 650-bp COX1 gene fragment generated only two major clusters among the 98 *Tor* spp. investigated in this study with very poor phylogenetic resolution. This result is consistent with that from Walton et al. (2017). The first cluster encompasses all *T. tambroides* fishes sampled from different regions of Indonesia, namely Bengkulu, Batang Tarusan River and Central Java. The second cluster contains the close clustering of *T. tambroides* with *T. tambra* from Malaysia and Indonesia. This is a clear indication that the standard COX1 gene fragment fails to distinguish *Tor* spp. even at the species level. The distinguishing power of this COX1 gene fragment seems less convincing, at least with regards to the heterogenous cluster encompassing fishes from three different origins such as Pahang and Terengganu states of Peninsular Malaysia, Sarawak state of East Malaysia as well as Java, Indonesia.

**Figure 2.**
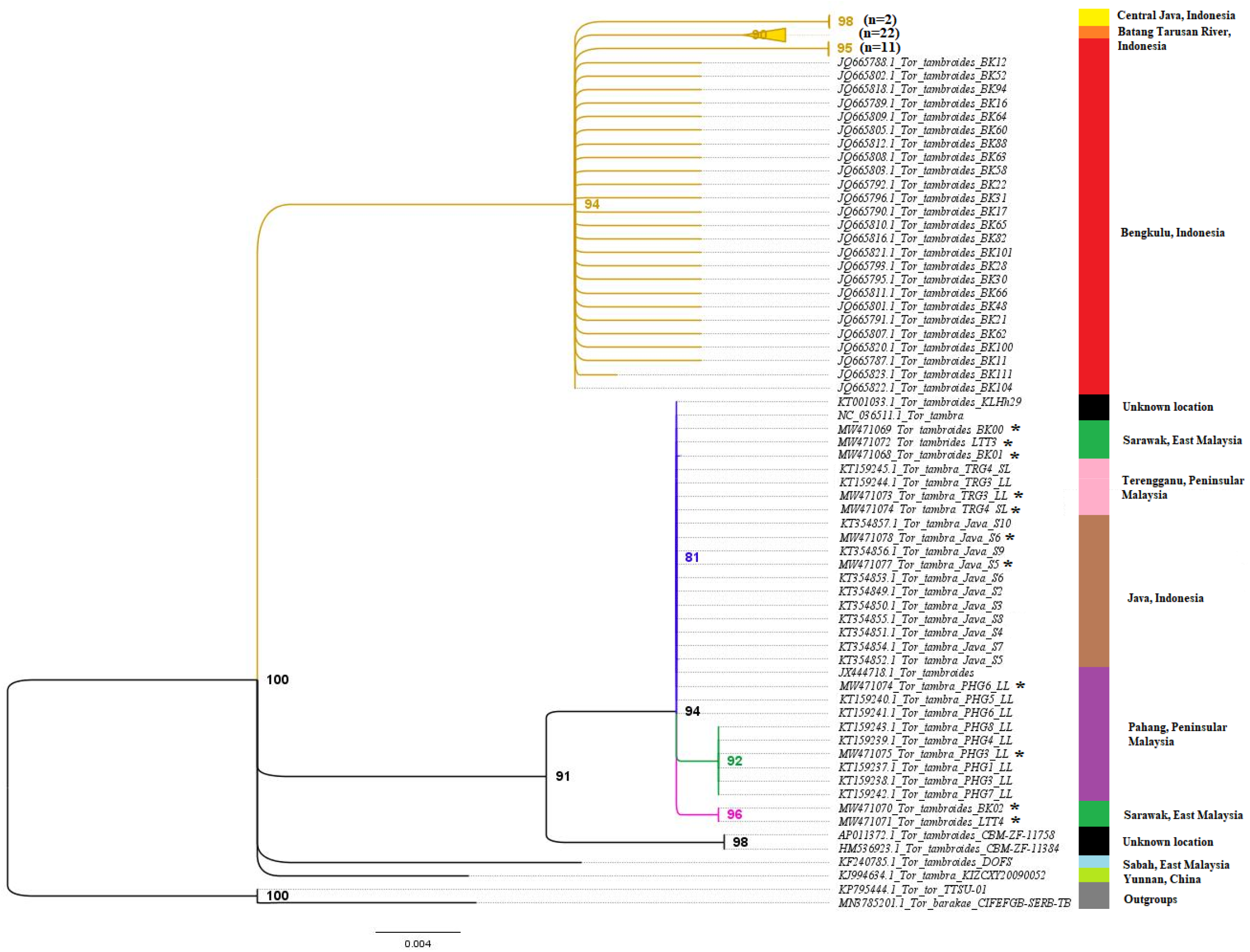
The maximum likelihood phylogenetic tree constructed based on the standard cytochrome oxidase I gene fragments of 98 *Tor* fishes with 1000 bootstrap replications. The sequences from this study were marked with an asterisk (*).

### Mitogenome-based phylogeny provides new insight into the population structure of Malaysian *Tor* spp

We utilized the 13 protein coding genes and two ribosomal RNA genes of all the 16 complete *Tor* mitogenomes (mainly *T. tambra* and *T. tambroides*) available from the GenBank database as well as from this study, to construct another tree in the attempt to improve the phylogenetic resolution and accuracy (Figure 3). At this taxon sampling level, we observed strongly supported monophyletic clustering based on sampling location for individuals from Peninsular Malaysia and Indonesia (Figure 3B) as compared to that constructed using the standard 650-bp COX1 gene fragments where most of them are clustered together and the phylogenetic relationship is poorly resolved (Figure 3A). On the contrary, three distinct mitolineages were observed among the Sarawak farm samples exhibiting paraphyletic relationships. The Sarawak state of Malaysia is one of the final remaining tropical forest sanctuaries on earth and this biodiversity hotspot is natural habitat to 18 million wildlife and indigenous people (Alamgir et al., 2020). Since the *Tor* fishes from Sarawak in this study were farmed fishes, it is essential to include wild *Tor* fishes from Sarawak in the future to comprehensively elucidate the geographical habitat origins of these Sarawak *T. tambroides* mitolineages. The sampling location of NC_036511.1 *T. tambra* fish is unknown but based on at its clustering with the other two Terengganu *T. tambra* fishes from this study, it is predicted that these three fishes may have sampled from the same locality. The *T. tambroides* CBM ZF 11758 (GenBank: AP011372.1) is genetically more divergent than all other *T. tambra* and *T. tambroides* fishes, with the lack of its sampling site record, it is postulated that either this fish has been misidentified as *T. tambroides* or possibly a cryptic species.

**Figure 3.**
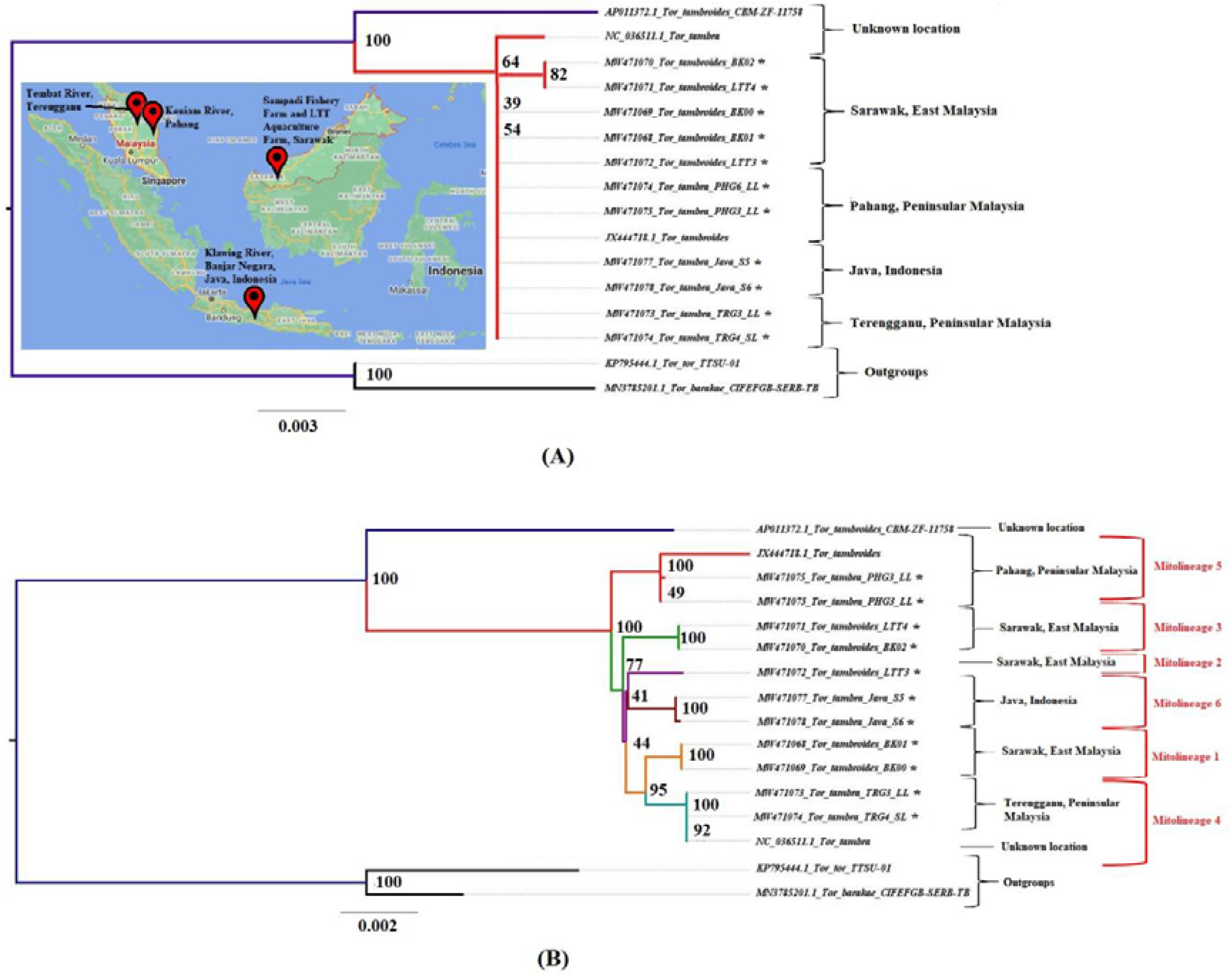
(A) The maximum likelihood phylogenetic tree constructed utilizing the 98 standard 650-bp COX1 gene fragments from 16 *Tor* fishes with 1000 bootstrap replications. The mitogenomes sequenced in this study were marked with an asterisk (*). The map shows the sampling sites of these *Tor* fishes. (B) The maximum likelihood phylogenetic tree constructed using 13 protein-coding genes and two ribosomal RNA genes across 16 *Tor* fishes, with 1000 bootstrap replications. The mitogenomes sequenced in this study were marked with an asterisk (*).

### Identification of population-specific mitochondrial gene markers among *Tor* spp. in Malaysia and Indonesia

Six mitolineages were identified in this study, which involved one population-specific mitolineage from Pahang, Terengganu and Java respectively as well as three Sarawak mitolineages (Table 2-6 & Supplementary Table 1-8). Among all the 15 mitochondrial genes examined, only five genes (namely COX1, Cytb, ND2, ND4 and ND5) contain SNPs that can differentiate the 6 putative populations based on the current taxon sampling. The complete COX1 and ND5 genes were the ones having the distinctively high amount of SNP markers with 21 and 19 markers detected respectively. Consistent with the genetic divergence of mitolineages from the Sarawak samples, several population-specific SNP markers were identified across these three mitolineages, suggesting that the current brood stock of farmed mahseer may have originated from three varying habitats and localities. Additional sampling of wild mahseers from this geographical region will be instructive to elucidate their mitogenome diversity and population structure. The mitolineage clade 5 grouped the two Pahang *T. tambra* from this study with the Pahang *T. tambroides* from Norfatimah et al. (2014), a similar phenomenon was observed in the phylogenetic tree. This suggests that there is possibility of cryptic *Tor* fishes present in the Keniam River of Pahang state. All the six population-specific mitolineage haplotypes identified in this study are powerful tools essential to distinguish biogeographically distinctive *Tor* populations.

**Table 2.** The population-specific SNP markers identified in the complete COX1 gene.

| Haplotypes for COX1 | Variable sites |  |  |  |  |  |  |  | Number of SNP marker |
| --- | --- | --- | --- | --- | --- | --- | --- | --- | --- |
|  | 3 | 4 | 7 | 7 | 9 | 1 | 1 | 1 |  |
|  | 3 | 4 | 4 | 5 | 9 | 0 | 1 | 2 |  |
|  |  | 4 | 7 | 6 | 2 | 1 | 9 | 4 |  |
|  |  |  |  |  |  | 1 | 1 | 5 |  |
| H1 (BK00 & BK01) | T | T | A | A | C | T | C | G | 2 |
| H2 (LTT3) | C | . | . | . | . | C | T | . | 1 |
| H3 (LTT4 & BK02) | C | C | . | . | T | C | . | . | 2 |
| H4 (TRG3_LL, TRG4_SL & NC_036511.1) | . | . | . | . | . | C | . | A | 1 |
| H5 (PHG3_LL, PHG6_LL & JX444718.1) | C | . | G | . | . | C | . | . | 1 |
| H6 (Java_S5 & Java_S6) | C | . | . | G | . | C | . | . | 1 |
| Total |  |  |  |  |  |  |  |  | 8 |

**Table 3.**
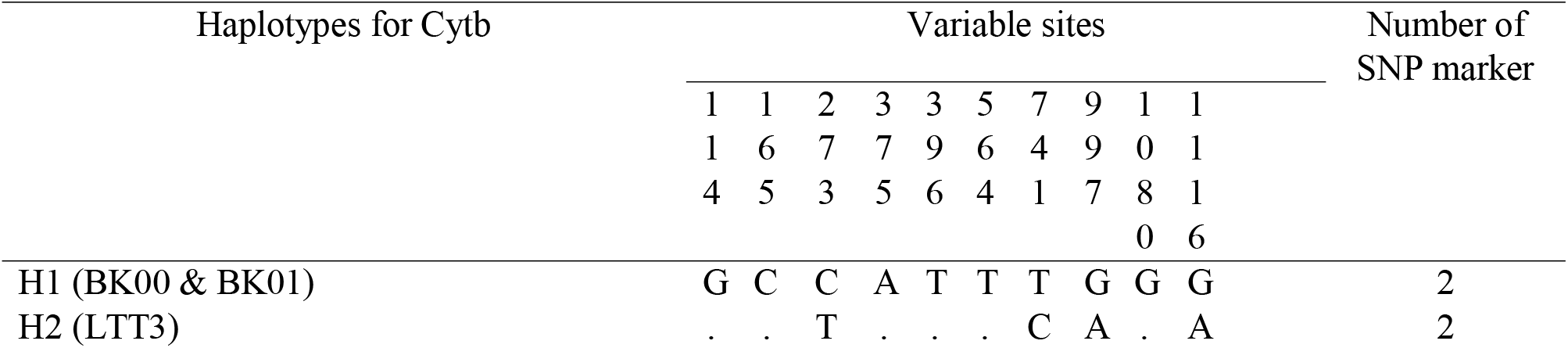
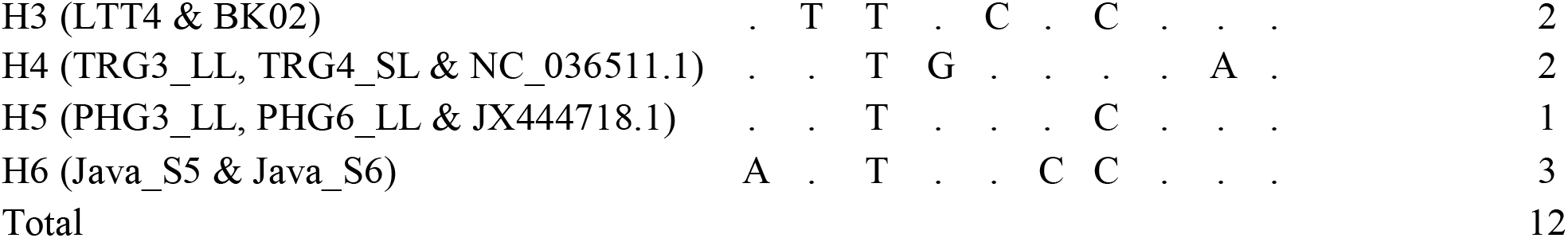
The population-specific SNP markers identified in the complete Cytb gene.

**Table 4.**
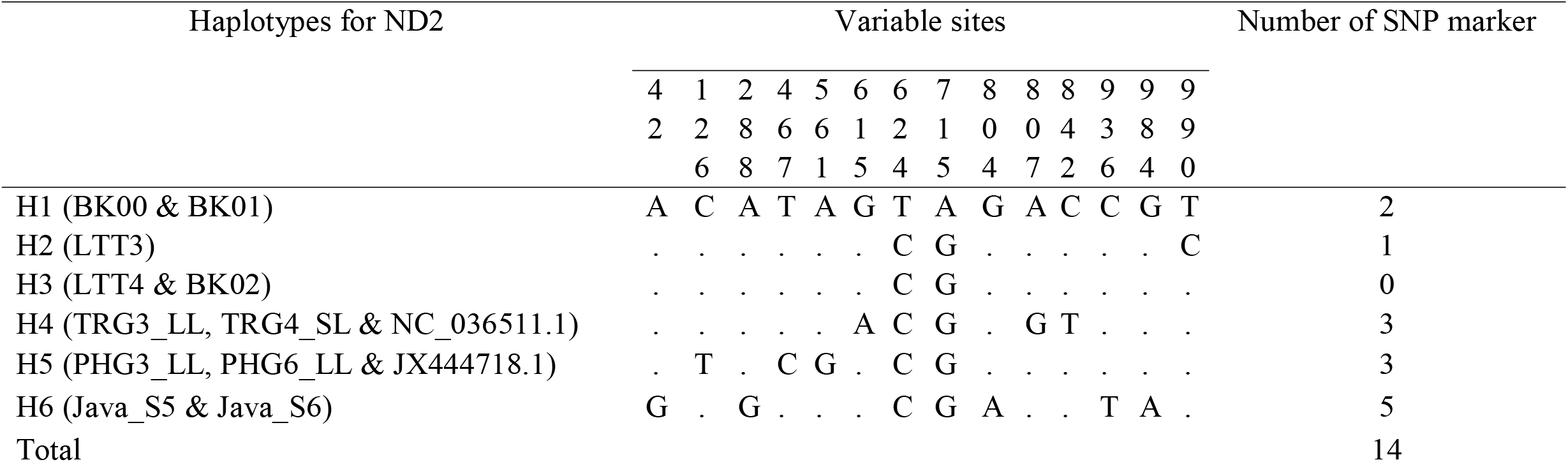
The population-specific SNP markers identified in the complete ND2 gene.

**Table 5.**
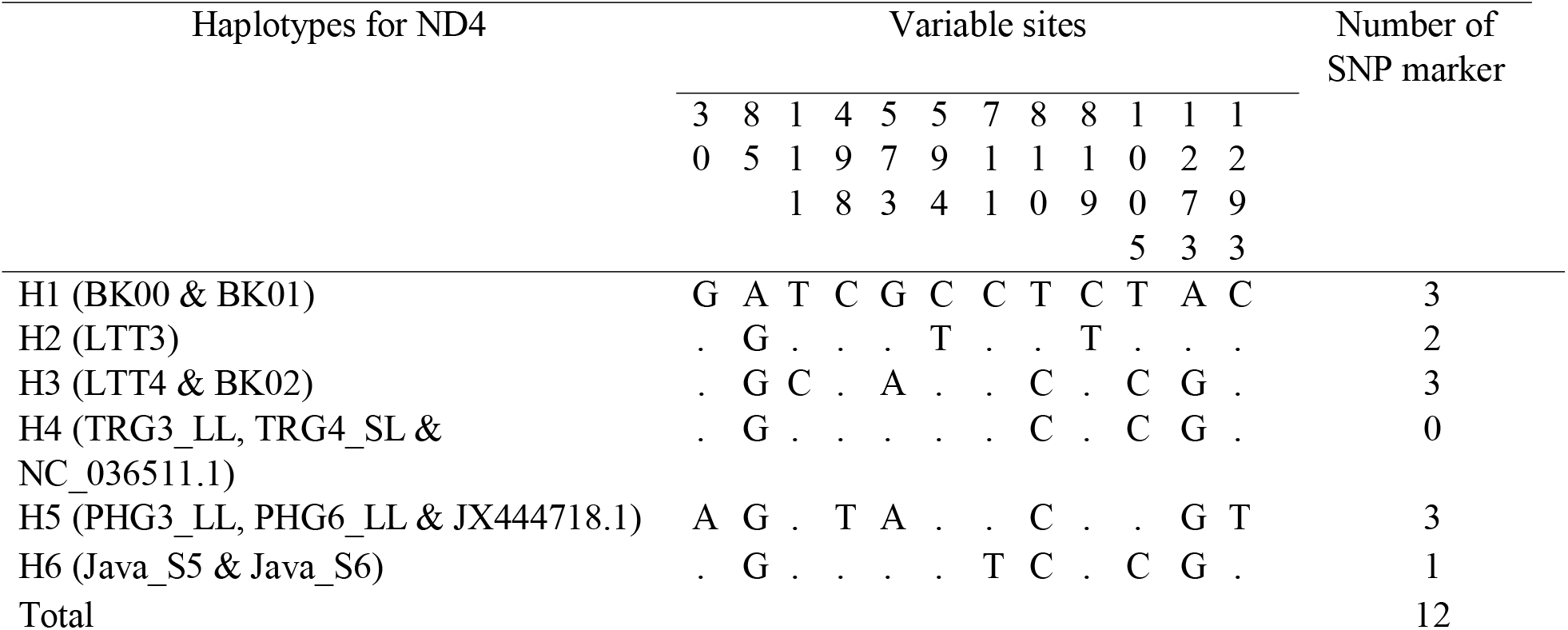
The population-specific SNP markers identified in the complete ND4 gene.

**Table 6.**
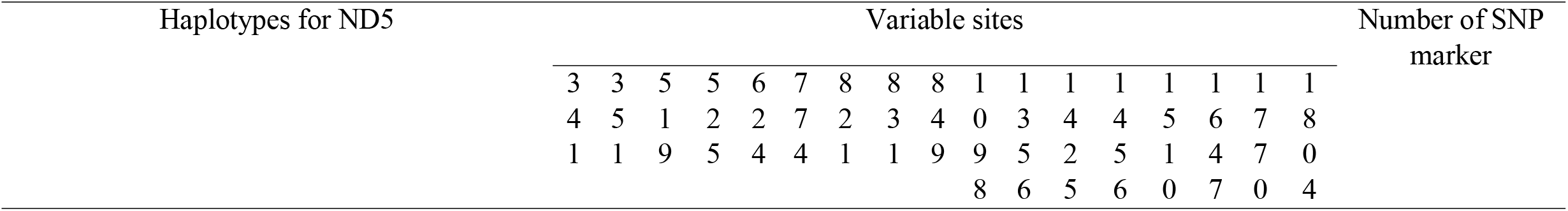
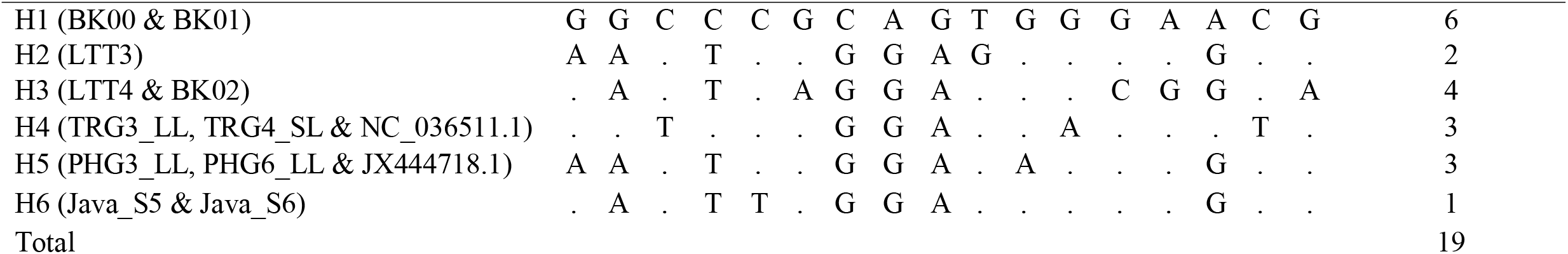
The population-specific SNP markers identified in the complete ND5 gene.

## Conclusion

In this study, 11 new *Tor tambroides* and *Tor tambra* complete mitogenomes from Java of Indonesia, Pahang and Terengganu of Peninsular Malaysia and Sarawak of Borneo Island, East Malaysia, were reported. The maximum likelihood phylogenetic tree was also constructed employing 15 genes (13 protein-coding and two ribosomal RNA gens) across the selected 16 *Tor* fishes, to improve the phylogenetic resolution plotted by utilizing the cytochrome oxidase I gene solely. Furthermore, five mitochondrial genes (namely COX1, Cytb, ND2, ND4 and ND5) contain population-specific SNP markers for all six *Tor* mitolineage haplotypes. However, given the lack of wild-caught specimen from Sarawak and its high mitogenome diversity, additional mitogenome sequencing of wild *Tor* spp from Sarawak and more generally the Borneo Island will be necessary to better inform the Mahseer broodstock management in the region.

## ACKNOWLEDGEMENT

This work was fully funded by Sarawak Research and Development Council through the Research Initiation Grant Scheme with grant number GL/F07/SRDC/03/2020 awarded to H. H. Chung.

---

**Supplementary Figure 1.**
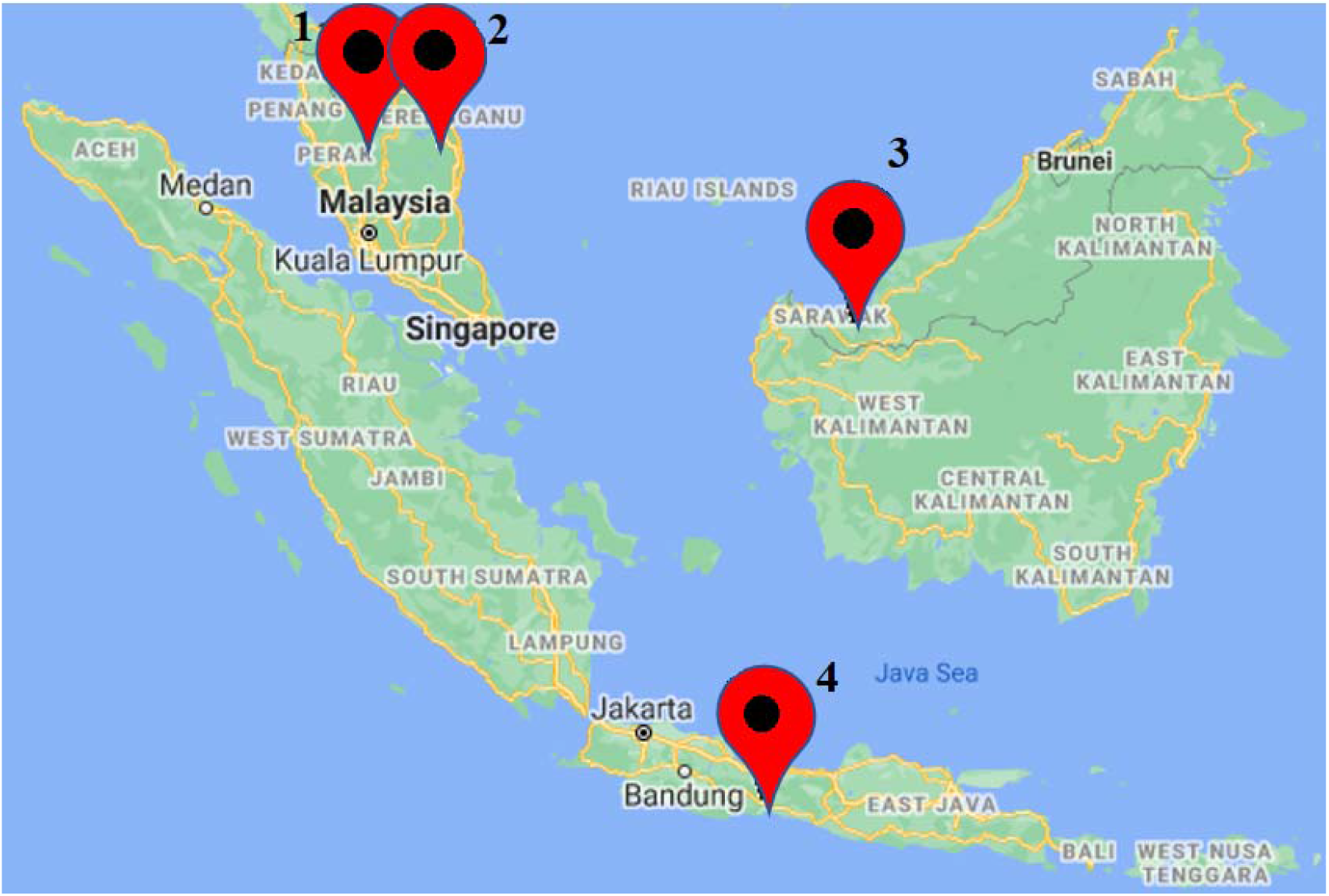
The sampling sites of *Tor* fishes in this study: 1. Keniam River, Pahang, Malaysia; 2. Tembat River, Terengganu, Malaysia; 3. Sampadi Fishery Farm and LTT Aquaculture Farm, Sarawak, Malaysia; 4. Klawing River, Banjar Negara, Java, Indonesia.

**Supplementary Figure 2.**
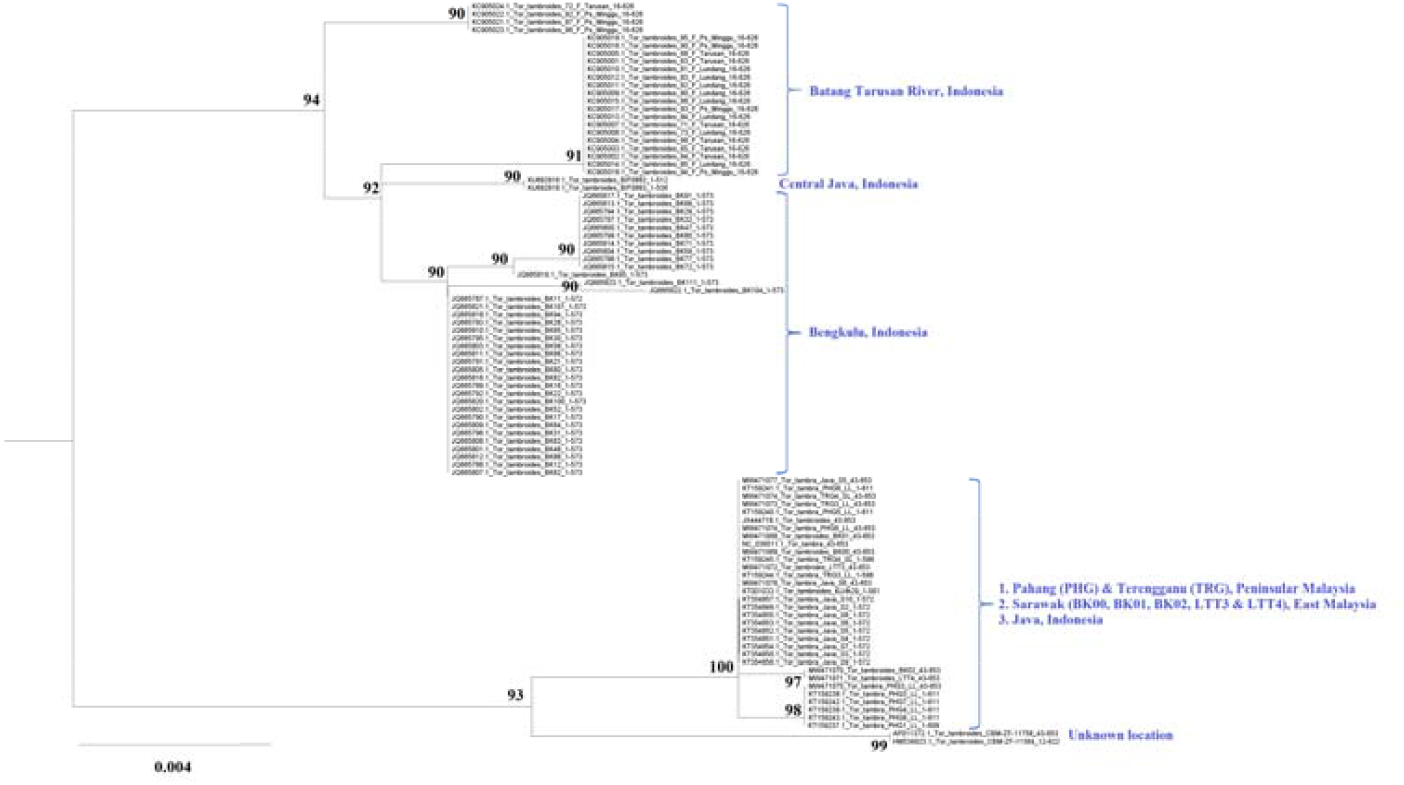
The maximum likelihood phylogenetic tree constructed based on the standard cytochrome oxidase I gene fragments of 98 *Tor* fishes with 1000 bootstrap replications.

**Supplementary Table 1.**
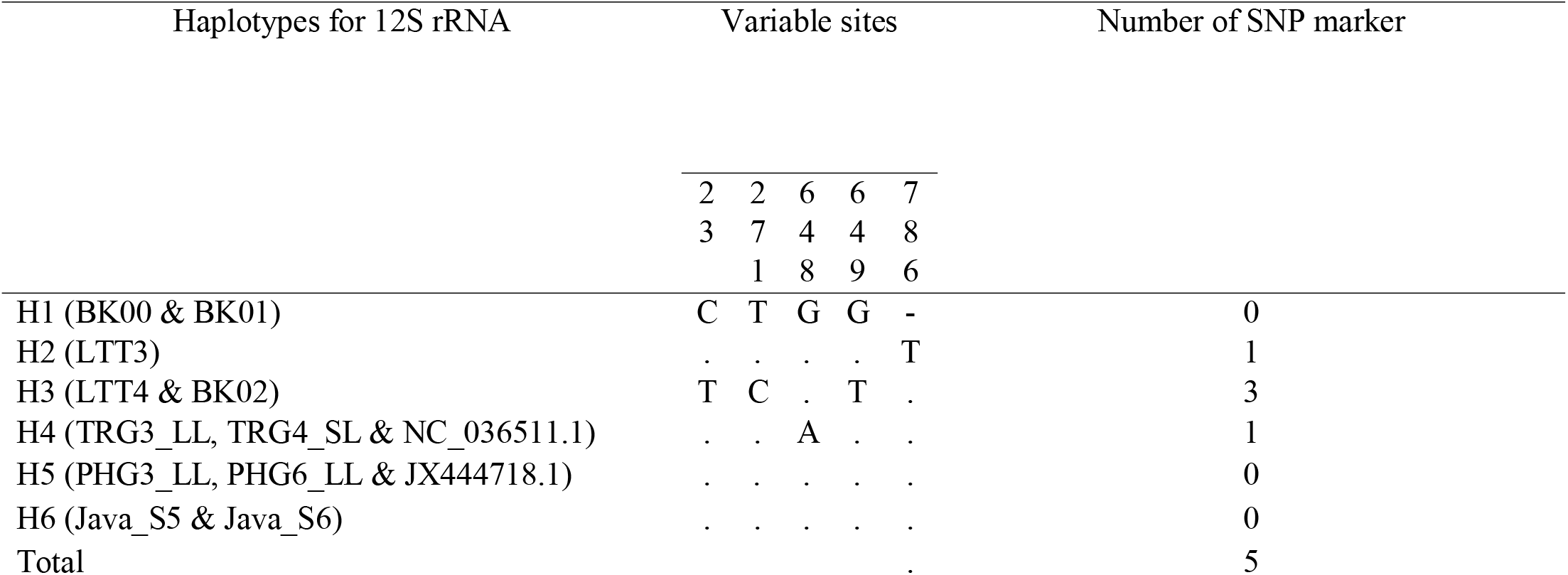
The population-specific SNP markers identified in the complete 12S rRNA gene.

| Haplotypes for 12S rRNA | Variable sites |  |  |  |  | Number of SNP marker |
| --- | --- | --- | --- | --- | --- | --- |
|  | 2 | 2 | 6 | 6 | 7 |  |
|  | 3 | 7 | 4 | 4 | 8 |  |
|  |  | 1 | 8 | 9 | 6 |  |
| H1 (BK00 & BK01) | C | T | G | G | - | 0 |
| H2 (LTT3) | . | . | . | . | T | 1 |
| H3 (LTT4 & BK02) | T | C | . | T | . | 3 |
| H4 (TRG3_LL, TRG4_SL & NC_036511.1) | . | . | A | . | . | 1 |
| H5 (PHG3_LL, PHG6_LL & JX444718.1) | . | . | . | . | . | 0 |
| H6 (Java_S5 & Java_S6) | . | . | . | . | . | 0 |
| Total |  |  |  |  | . | 5 |

**Supplementary Table 2.**
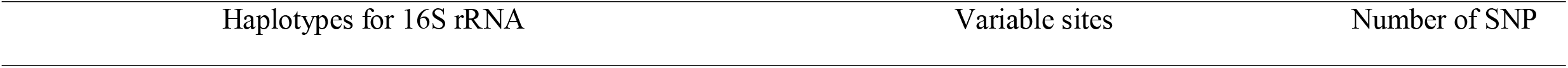
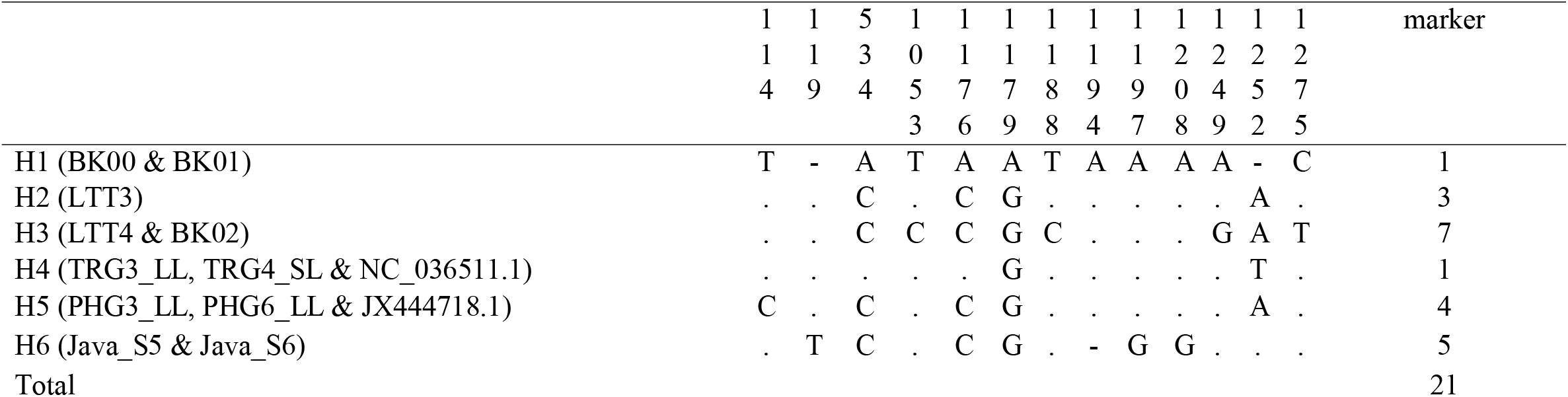
The population-specific SNP markers identified in the complete 16S rRNA gene.

| Haplotypes for 16S rRNA | Variable sites | Number of SNP |
| --- | --- | --- |

|  |  |  |  |  |  |  |  |  |  |  |  |  |  |  |
| --- | --- | --- | --- | --- | --- | --- | --- | --- | --- | --- | --- | --- | --- | --- |
|  | 1 | 1 | 5 | 1 | 1 | 1 | 1 | 1 | 1 | 1 | 1 | 1 | 1 | marker |
|  | 1 | 1 | 3 | 0 | 1 | 1 | 1 | 1 | 1 | 2 | 2 | 2 | 2 |  |
|  | 4 | 9 | 4 | 5 | 7 | 7 | 8 | 9 | 9 | 0 | 4 | 5 | 7 |  |
|  |  |  |  | 3 | 6 | 9 | 8 | 4 | 7 | 8 | 9 | 2 | 5 |  |
| H1 (BK00 & BK01) | T | - | A | T | A | A | T | A | A | A | A | - | C | 1 |
| H2 (LTT3) | . | . | C | . | C | G | . | . | . | . | . | A | . | 3 |
| H3 (LTT4 & BK02) | . | . | C | C | C | G | C | . | . | . | G | A | T | 7 |
| H4 (TRG3_LL, TRG4_SL & NC_036511.1) | . | . | . | . | . | G | . | . | . | . | . | T | . | 1 |
| H5 (PHG3_LL, PHG6_LL & JX444718.1) | C | . | C | . | C | G | . | . | . | . | . | A | . | 4 |
| H6 (Java_S5 & Java_S6) | . | T | C | . | C | G | . | - | G | G | . | . | . | 5 |
| Total |  |  |  |  |  |  |  |  |  |  |  |  |  | 21 |

**Supplementary Table 3.** The population-specific SNP markers identified in the complete ATPase 6 gene.

| Haplotypes for ATPase 6 | Variable sites |  |  |  |  | Number of SNP marker |
| --- | --- | --- | --- | --- | --- | --- |
|  | 4 | 1 | 1 | 2 | 6 |  |
|  | 8 | 4 | 8 | 9 | 6 |  |
|  |  | 4 | 0 | 4 | 0 |  |
| H1 (BK00 & BK01) | T | G | G | A | T | 0 |
| H2 (LTT3) | . | . | . | . | C | 1 |
| H3 (LTT4 & BK02) | . | . | A | . | . | 1 |
| H4 (TRG3_LL, TRG4_SL & NC_036511.1) | C | . | . | . | . | 1 |
| H5 (PHG3_LL, PHG6_LL & JX444718.1) | . | A | . | G | . | 2 |
| H6 (Java_S5 & Java_S6) | . | . | . | . | . | 0 |
| Total |  |  |  |  |  | 5 |

**Supplementary Table 4.** The population-specific SNP markers identified in the complete COX2 gene.

| Haplotypes for COX2 | Variable sites | Number of<br>SNP marker |
| --- | --- | --- |
|  | 5<br>2<br>2 |  |
| H1 (BK00 & BK01) | G | 0 |
| H2 (LTT3) | . | 0 |
| H3 (LTT4 & BK02) | . | 0 |
| H4 (TRG3_LL, TRG4_SL & NC_036511.1) | . | 0 |
| H5 (PHG3_LL, PHG6_LL & JX444718.1) | . | 0 |
| H6 (Java_S5 & Java_S6) | A | 1 |
| Total |  | 1 |

**Supplementary Table 5.**
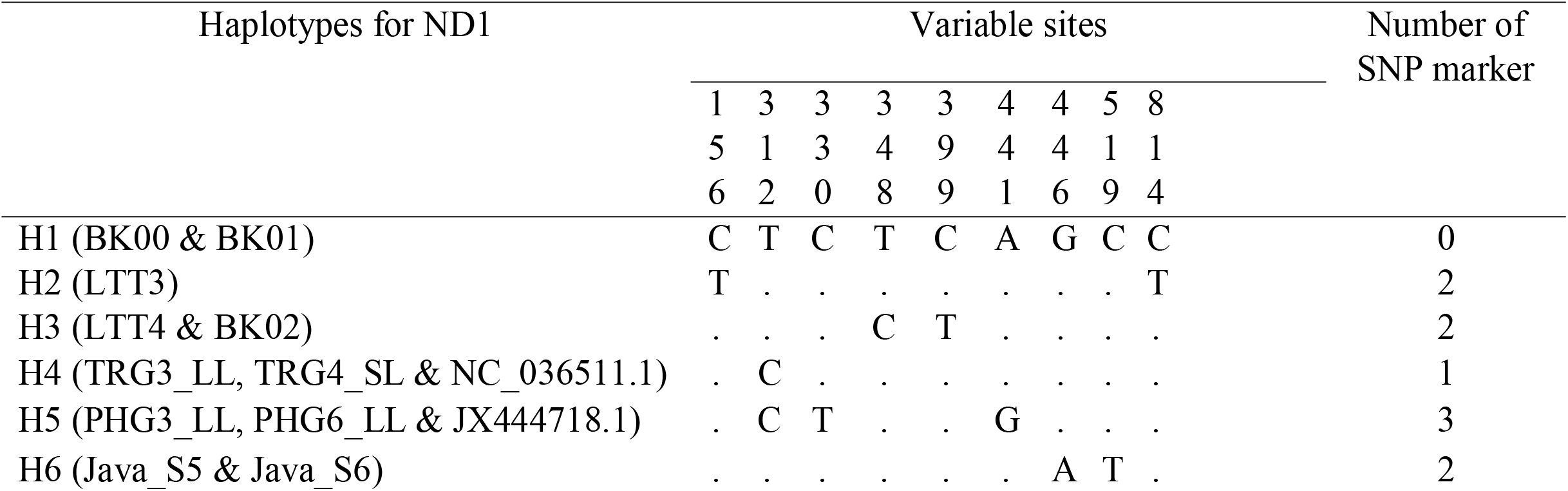
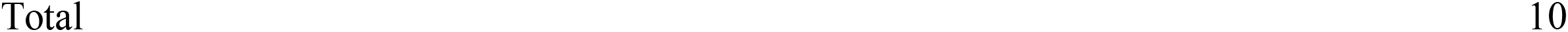
The population-specific SNP markers identified in the complete ND1 gene.

**Supplementary Table 6.** The population-specific SNP markers identified in the complete COX3 gene.

| Haplotypes for COX3 | Variable sites |  |  |  |  |  | Number of<br>SNP marker |
| --- | --- | --- | --- | --- | --- | --- | --- |
|  | 2 | 4 | 5 | 5 | 5 | 7 |  |
|  | 4 | 0 | 7 | 8 | 9 | 1 |  |
|  | 0 | 5 | 6 | 8 | 1 | 1 |  |
| H1 (BK00 & BK01) | A | C | T | G | C | T | 0 |
| H2 (LTT3) | . | . | . | . | . | . | 0 |
| H3 (LTT4 & BK02) | . | . | . | A | . | . | 1 |
| H4 (TRG3_LL, TRG4_SL & NC_036511.1) | G | . | . | A | . | . | 2 |
| H5 (PHG3_LL, PHG6_LL & JX444718.1) | . | T | . | . | . | C | 2 |
| H6 (Java_S5 & Java_S6) | . | . | C | . | T | . | 2 |
| Total |  |  |  |  |  |  | 7 |

**Supplementary Table 7.**
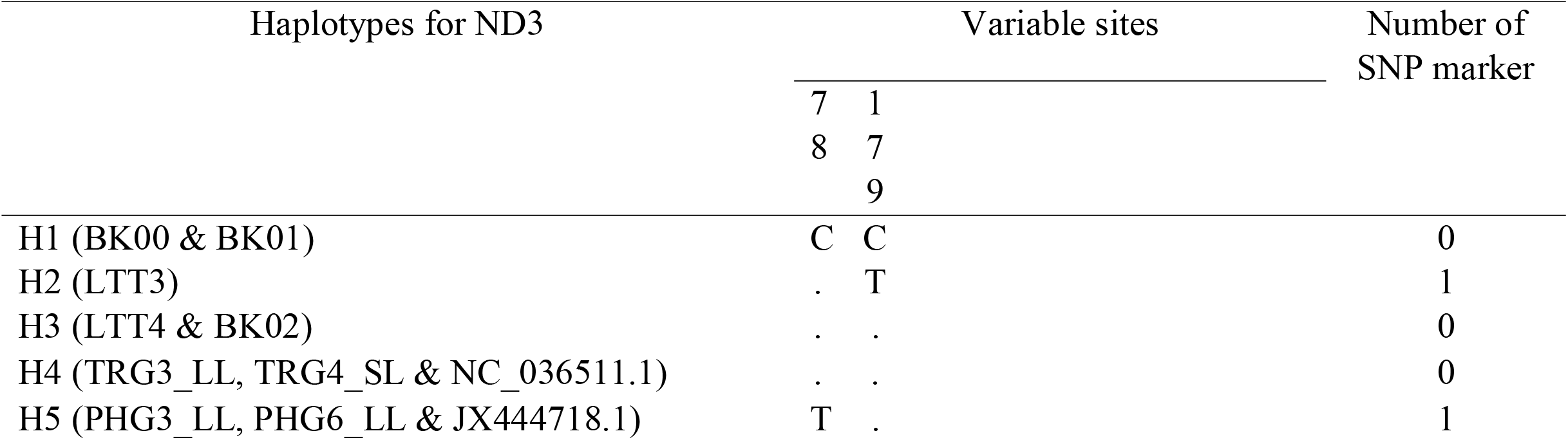
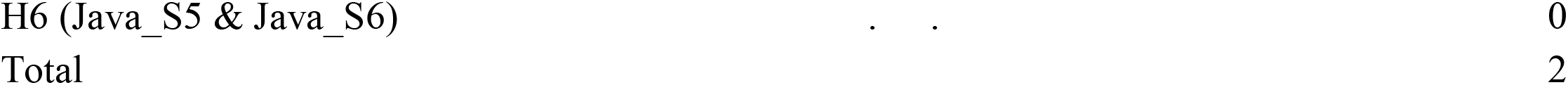
The population-specific SNP markers identified in the complete ND3 gene.

**Supplementary Table 8.** The population-specific SNP markers identified in the complete ND6 gene.

| Haplotypes for ND6 | Variable sites |  |  |  | Number of<br>SNP marker |
| --- | --- | --- | --- | --- | --- |
|  | 2 | 3 | 3 | 3 |  |
|  | 8 | 1 | 3 | 9 |  |
|  |  | 8 | 6 | 6 |  |
| H1 (BK00 & BK01) | A | A | C | T | 1 |
| H2 (LTT3) | . | G | . | C | 1 |
| H3 (LTT4 & BK02) | . | . | . | C | 0 |
| H4 (TRG3_LL, TRG4_SL & NC_036511.1) | . | . | . | C | 0 |
| H5 (PHG3_LL, PHG6_LL & JX444718.1) | G | . | T | C | 2 |
| H6 (Java_S5 & Java_S6) | . | G | . | C | 1 |
| Total |  |  |  |  | 5 |

